# Experienced Social Partners Hinder Learning Performance in Naive Clonal Fish

**DOI:** 10.1101/2022.10.13.512085

**Authors:** Fritz A. Francisco, Almond Stöcker, Juliane Lukas, Pawel Romanczuk, David Bierbach

## Abstract

Social learning is widely assumed to enhance individual learning efficiency, particularly when naive observers have access to skilled demonstrators, yet the conditions under which this assumption holds remain poorly understood. Here, we investigated how individual learning is influenced by the skill level of social partners. We predicted that naive individuals would benefit from observing experienced conspecifics, yet found the opposite: trained partners significantly impaired naive individuals’ learning performance, while trained individuals remained unaffected by their partner’s skill level. We conducted experiments in near-identical individuals, using the all-female clonal Amazon molly (*Poecilia formosa*) to test whether these fish can learn an operant foraging task, whether individuals differ consistently in learning ability, and whether partner skill level influences learning performance. Using an operant conditioning paradigm over five days, half of the fish were trained to locate food inside a cylinder, whereas the remaining individuals received food randomly dispersed within their tank. Trained individuals subsequently visited the cylinder more frequently than randomly fed individuals and exhibited consistent individual differences in learning performance. In a second phase, fish were allowed to observe a conspecific while individual training was either continued (for trained individuals) or initiated (for naive individuals). We found that trained individuals did not benefit from the presence of a partner, regardless of the partner’s proficiency, but consistently outperformed naive individuals. In contrast, naive individuals showed reduced learning performance when paired with experienced partners, but not with naive partners. Together, our results indicate that Amazon mollies achieve this foraging task through individual learning and exhibit stable differences in learning ability. Moreover, social learning depends on both the learner’s own skill level and that of the social partner, such that observing an experienced conspecific may, in some cases, impede rather than enhance learning.

## Introduction

As early as 1514, Machiavelli observed that “Men nearly always follow the tracks made by others and proceed in their affairs by imitation, even though they cannot entirely keep to the tracks of others or emulate the prowess of their models.” This tendency is not unique to humans: many gregarious animals acquire information about their environment from social partners [1, 2]. Such information use is commonly referred to as observational or social learning [3–5]. Social learning contrasts with private learning, in which information is gained through individual experience in the absence of others [6, 7].

While private learning relies on direct interaction with the environment, social learning involves observing others and copying their actions [8]. Social learning can be particularly advantageous when individuals differ in their knowledge of the environment. For example, task-naive Amazon parrots (*Amazona amazonica*) copy the behaviour of more experienced individuals to access an obstructed food source [9], and wild guppies (*Poecilia reticulata*) copy both food-patch preferences and predator-avoidance behaviour from conspecifics [10]. More broadly, the presence of conspecifics can lower predation risk [11] and improve foraging efficiency [12], and social learning allows individuals to bypass extensive individual exploration, thereby reducing time, energy expenditure, and opportunity costs [4, 10, 13–18].

However, these benefits are not necessarily shared equally by all group members [19, 20], and the presence of a social partner can also impose costs. Early work examining mismatch and costs in observer–demonstrator relationships laid conceptual foundations for this [21–23], on which the present study builds. Such costs may arise from distraction [17] or from altered time budgets that favour social interactions over task engagement [24]. Importantly, demonstrator skill alone does not determine the value of social information: in guppies, familiarity between observer and demonstrator appears to outweigh demonstrator proficiency in shaping social learning [25]. Observed individuals can differ substantially in performance and, consequently, in the quality of information they provide.

In a discrimination learning task in pigeons, for instance, Biederman and Vanayan [22] showed that individuals who observed others making mistakes during learning outperformed those who were shown stereotypic and proficient solutions. Similarly, in maze-solving tasks, Swaney et al. [13] showed that experienced individuals complete the task rapidly but provide naive followers little opportunity to learn the route themselves, while less skilled demonstrators were followed more readily than proficient ones. Together, these findings suggest that the value of social information depends not only on availability but on its usability, giving rise to a trade-off between private and social information that is shaped by demonstrator behavior as much as by demonstrator experience [25–27]. Individual exploration and private information acquisition may therefore be favoured not only when social partners possess equivalent prior experience, rendering social information redundant [7, 28], but also when experienced demonstrators fail to provide accessible or transferable information, such that demonstrator skill does not always translate into a usable signal for the observer [13, 22].

Yet despite evidence that demonstrator experience does not straightforwardly translate into observer benefit, the assumption that more skilled partners invariably benefit naive observers has rarely been tested [13, 29, 30], and the possibility that unequal skill levels and information asymmetry between partners could hinder rather than facilitate learning remains poorly understood. The capacity to benefit from social information may itself depend on the learner’s own cognitive characteristics, including their inherent learning ability. Individuals within a population often differ consistently in behavioral and cognitive traits, including learning ability [31, 32]. Such among-individual variation has been documented even in genetically identical organisms [33, 34], suggesting that factors beyond genetic composition shape individual phenotypes. Whether learning ability itself varies consistently among individuals, and how such variation interacts with social context, remains an open question.

In this study, we investigated how variation in skill level between genetically near-identical social partners influences learning in naive individuals and overall task performance in experienced individuals. We used the Amazon molly (*Poecilia formosa*), a naturally occurring clonal fish species native to the border region of the United States and Mexico [35, 36]. These all-female fish reproduce gynogenetically, producing live offspring genetically identical to their mothers and sisters [37–39]. Owing to its clonal genetic background and gregarious lifestyle, this species has been proposed as a powerful model for studying consistent individual behavioural differences and their effects on group-level dynamics [40–43]. Using genetically identical individuals minimises the contribution of genetic differences to variation in social learning outcomes, allowing experiential factors to be examined more directly [44]. However, learning abilities in this species have received limited attention [45].

All individuals used in this study shared an near-identical genetic composition and highly similar rearing conditions. During an initial phase of private information acquisition, we employed an operant conditioning paradigm over five days (six training sessions per day) to generate two cohorts differing in experience. One cohort was trained to locate food inside a white, opaque cylinder with a ceramic base (the task; Fig. 1), thereby acquiring task experience. The second cohort received food randomly distributed throughout the tank, preventing the formation of an association between food location and the cylinder and thus remaining task-naive. In a subsequent phase of social information acquisition, pairs of individuals were given visual access to one another, enabling mutual observation while training continued for experienced fish and commenced for naive fish (five days, six sessions per day). This fully factorial design yielded all possible experience combinations: naive–naive, naive–trained, and trained–trained pairs.

**Figure 1.**
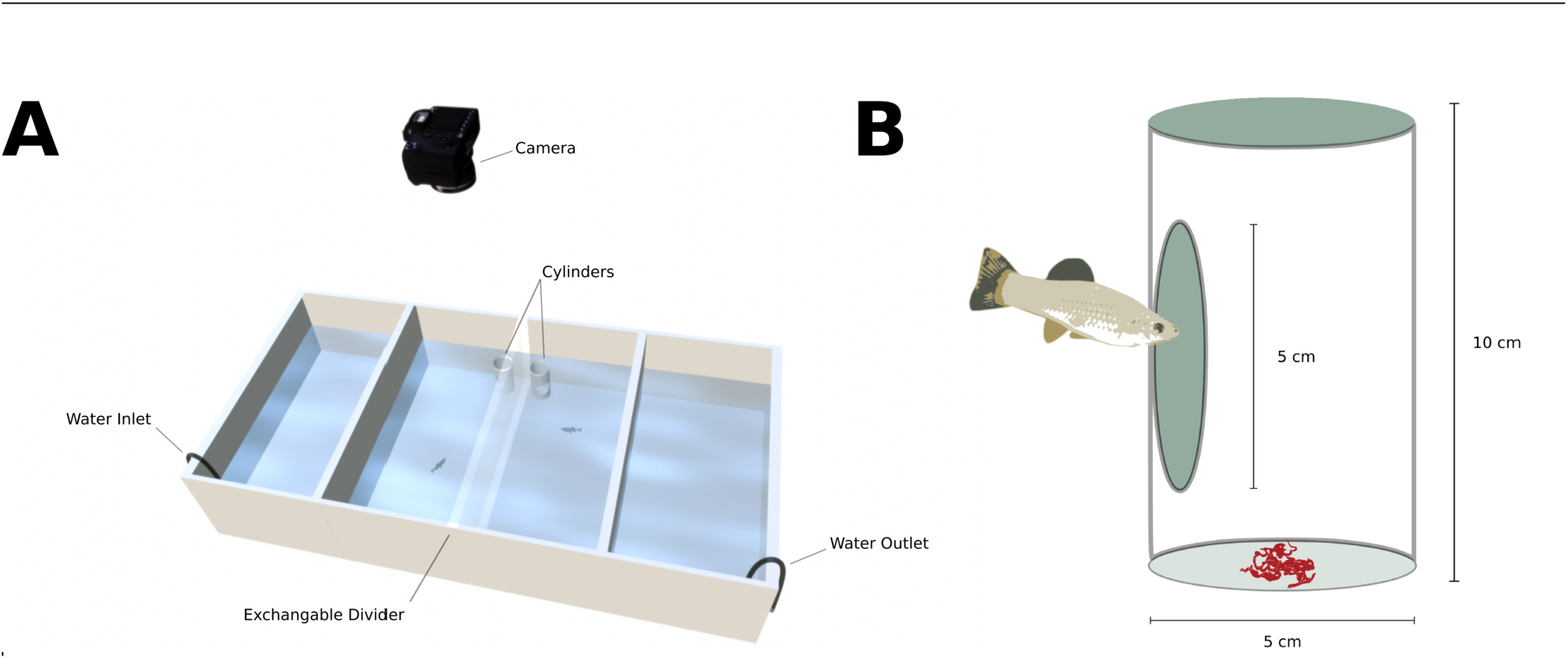
**A** Schematic of the general recording setup. Each inlet and outlet was attached to an individual circulating filter system. **B** Cylinder serving as concealed food source used in the conditioning trials. Food was presented within an opaque cylinder, that could only be accessed through a horizontal opening. Entry into the cylinder was monitored through the top opening, vertically facing the camera. The cylinders were glued to ceramic plates to ensure stability. This further ensured that food particles and olfactory cues were contained within the cylinder.

Using this design, we tested three hypotheses. Hypothesis I was that Amazon mollies can learn the task; we predicted that trained individuals would show significantly increased visit rates to the cylinder over successive training sessions relative to naive individuals (P1). Hypothesis II was that individuals exhibit consistent differences in learning performance; we predicted that learning rates among trained fish would show significant inter-individual variance (P2). Hypothesis III proposed that partner skill level affects learning and overall performance once social information becomes available. From this hypothesis we derived two predictions: first, that naive fish paired with trained partners would be more likely to access the food source than those paired with other naive individuals (P3); and second, that experienced fish would show reduced task performance when paired with naive partners relative to similarly experienced partners (P4). These predictions are not symmetric, as the first concerns information gain by naive observers, while the second concerns potential costs to experienced individuals, and therefore rest on distinct but complementary mechanisms.

## Materials and Methods

### Study Organism and Maintenance

We used the Amazon molly (*Poecilia formosa*), a naturally occurring clonal freshwater fish, as our study organism. This all-female species originated approximately 100,000 years ago from a rare hybridization event between a male sailfin molly (*P. latipinna*, ♂) and a female Atlantic molly (*P. mexicana*, ♀) [37–39, 46–48]. The Amazon molly reproduces via gynogenesis, whereby females require sperm from males of closely related poeciliid species to initiate embryogenesis [49]. However, no paternal genetic material is incorporated, resulting in broods of offspring that are genetically identical to one another and to their mothers [50]. The clonal lineage used in this study has been maintained in captivity for many generations. Fish were bred using *P. mexicana* males as sperm donors at the animal care facilities of Humboldt-Universität zu Berlin. Fish were housed in 200-L tanks filled with aged tap water and maintained at 26 °C. They were fed twice daily *ad libitum* with commercially available flake food (Sera Vipan Baby Nature) and thawed bloodworms (*Chironomidae* sp.). All experimental procedures were approved by the German State Office for Health and Social Affairs (LAGeSo; animal experiment number #0089/21).

### Experimental Design

The experiment consisted of two phases. In the first phase, we established two treatment groups. One group was fed exclusively inside an opaque cylinder three times per day for one week (hereafter, “trained”; Figure 1). The second group received food randomly dispersed throughout the experimental tank and thus had no opportunity to associate food with the cylinder (hereafter, “naive”). In the second phase, fish were visually paired either with an individual from the same or from a different training regime. Feeding in the cylinder was continued for previously trained individuals and initiated for naive individuals. Each experimental tank (300 × 600 × 200 mm) was divided into two equal-sized compartments by an opaque partition, with one fish housed in each compartment. During experimental trials, the opaque divider could be replaced with a transparent one to allow visual interaction between paired individuals (Figure 2). All tanks were externally filtered (EHEIM Professional III 250) throughout the experiment to maintain water quality and permit the exchange of olfactory cues. Water quality (pH, NH_3_/NH_4_, NO_2_, NO_3_; SERA drip test kits) was monitored weekly, and 50% of the water was replaced at the same interval. Water temperature was maintained between 23 and 26 °C via ambient room temperature. Water depth was kept at 70 mm, resulting in a total volume of 18.7 L per tank and 3.5 L per compartment. To facilitate learning, fish were maintained under continuous illumination, a regime shown to improve learning performance without increasing stress in a closely related species [51]. All fish were fed frozen bloodworms, prepared to thaw approximately 30 min before each experimental session

**Figure 2.**
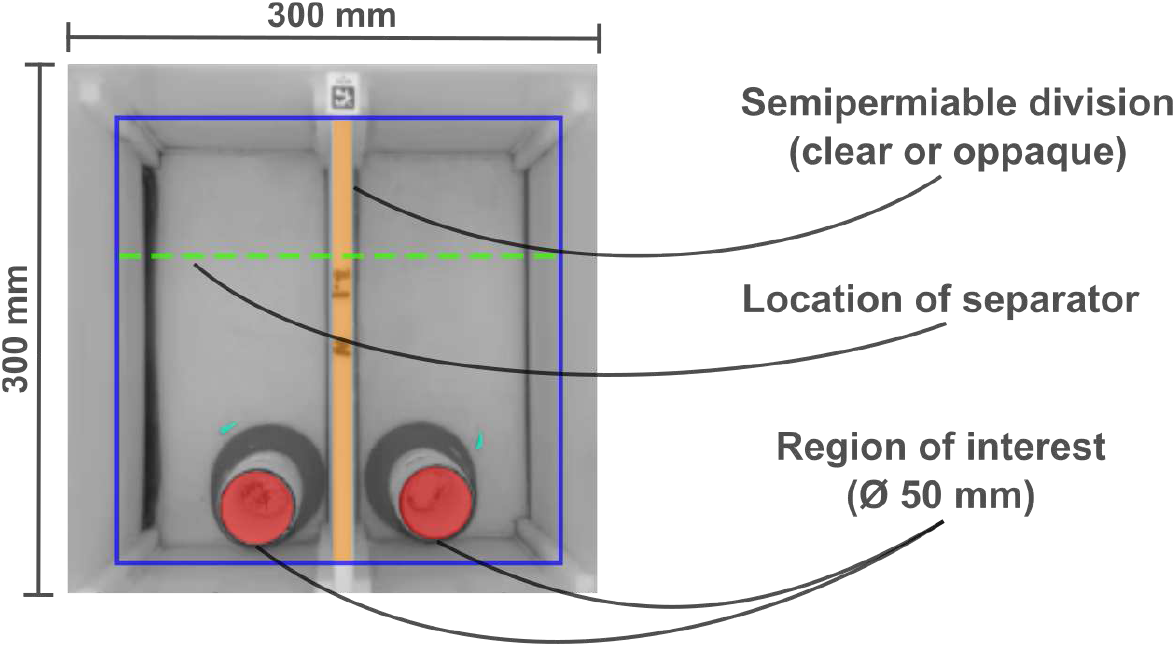
Recorded view of an experimental tank, showing the centrally located compartments housing one individual on each side (light blue). The location at which the cylinder serving as the food source was placed is denoted as the region of interest (red). The exchangeable divider, which could be either clear or opaque, is shown in the middle (orange). The separator used to standardize starting distance in each test instance is shown as a dashed line (green). The water level is highlighted along the tank edges (blue).

### Food Conditioning Experiments

#### Private Information — Week 1

At the start of the experiment, pairs of individuals were created (N_Total_ = 36, N_Pairs_ = 18) and randomly assigned to one of six experimental tanks, with each individual randomly assigned to one of two separate compartments.

Fish were sourced from multiple husbandry tanks to avoid familiarity among individuals and size-matched (standard length = 23 ±2mm) to control for metabolic needs, swimming ability, and age differences [52, 53]. For individual conditioning, individuals were randomly designated to be either trained or naive within each of six simultaneously trained pairs (N_trained_=18, N_naive_=18). Trained individuals were trained six times per day for five days without visual access to their conspecific partner. Naive individuals received food but were not trained on the foraging task. Each training session lasted eight minutes and was recorded using consumer-grade webcams (Logitech C920 HD Pro, USB 3.0; 432 ×240px; grayscale; 30 frame per second; see Supporting Material, Video Recording Script) mounted above each individual compartment. Cameras were centered above the tanks to minimize occlusions and perspective distortion and to ensure that any residual distortion was evenly distributed across individuals. During each training session, individuals were either presented with an opaque, vertically oriented PVC cylinder (height: 100 mm, diameter: 50 mm; Figure 1) containing food (true stimulus) or underwent a mock trial. The order of true and mock trials was randomized daily while maintaining equal representation (3 true and 3 mock trials per day). In mock trials, all procedural steps were identical, including confinement and subsequent removal of the separator, but no stimulus or food was presented. Mock trials were included to reduce associations with procedural cues and to ensure that learning was specific to the task stimulus [54–56]. To standardize the initial distance to the food source, individuals were confined to one side of the compartment at the start of each session using a temporary separator (Figure 2).

For individuals in the trained cohort, the cylinder contained approximately eight bloodworms that were visually occluded and accessible only through a circular opening on the side of the cylinder (Figure 1). Individuals in the naive cohort experienced identical conditions, except that food was distributed randomly throughout the compartment and remained accessible for the duration of the session. At the end of each session, the cylinder and any remaining food particles were removed using a pipette.

#### Social Information — Week 2

In the second week of the experiment, individuals were reassigned to new size-matched partners and randomly redistributed across the six experimental tanks to ensure that each fish experienced a novel testing environment. Based on each individual’s prior training status and that of its partner, four social treatments were created: trained–trained (TT; N=12), naive–trained (NT; N=6), trained–naive (TN; N=6), and naive–naive (NN; N=12). The uneven number of individuals across treatments is due to the pairing and symmetrical design in trained- trained and naive-naive. The opaque divider separating the two compartments was replaced with a transparent partition, allowing full visual access between paired individuals (Figure 2). This divider remained in place throughout the five consecutive days of the social phase. During this period, all individuals were trained and tested following the same individual conditioning protocol described above (Figure 1).

### Video Analysis

To quantify learning outcomes, fish were tracked using a custom Python-based tracking pipeline (`track2h5()`; see Supporting Material, Tracking Code) implemented with the computer vision library OpenCV [57]. Fish were detected through background subtraction and tracked over subsequent frames. Detected objects were filtered by size and speed and further refined using an isolation forest algorithm to restrict detections to actual fish and minimize noise from reflections and moving particles, such as air bubbles.

Individual positions were recorded as two-dimensional Cartesian coordinates corresponding to the center of mass of each filtered detection contour. Because background subtraction can fail when animals remain stationary, missing positions were linearly interpolated over time. The first 10 s of each test instance were considered an acclimation phase following removal of the separator and were excluded from analysis. To standardize recordings, all videos were truncated to a maximum duration of 480 s (8 min.), resulting in an analyzed period of 470 s per instance.

Because individuals were confined to fixed compartments, identities were maintained based on spatial location.

The position of the stimulus cylinder was detected automatically using a Hough transform from the OpenCV library, yielding the center coordinates and radius of the cylinder [57]. This enabled precise calculation of the Euclidean distance between each individual and the cylinder center at each time point. All videos were additionally inspected manually to verify the accuracy of cylinder detection and tracking results.

### Statistical Analysis

All statistical analyses were conducted in R (version 3.6.3, Holding the Windsock). Statistical inference was based on generalized linear mixed-effects models (logit models) fitted using the `glmer` function from the `lme4` package. Model selection was performed using Akaike’s Information Criterion (AIC) for fixed effects, or using conditional AIC (cAIC) for random effects, implemented via the `cAIC4` package [58]. Model diagnostics and accuracy assessments were carried out using `check_model` from the `performance` package [59], and test statistics were summarized using `tab_model` from the `sjPlot` package. Variance components were evaluated using the boundary correction method described by Stram and Lee [60], which was originally proposed for linear mixed models but is also applicable in the generalized case [61]. Statistical significance was assessed at the 95% level, and all confidence intervals (CIs) are reported as 95% CIs.

Individuals (*i* = 1, … , 36), were defined as having reached the region of interest (i.e. solved the task) during test instance *j* = 1, … , 15 if their distance to the center of the cylinder was less than 2.5 cm for at least 1 s. This outcome was coded as *y*_*ij*_ = 1; failure to meet this criterion was coded as *y*_*ij*_ = 0.

Assuming that repeated feeding inside the cylinder increases the likelihood of solving the task, individual learning performance was modeled as the probability of reaching the region of interest using a logistic regression framework. Two related model variants were used to analyze data from Week 1 (Model 1) and Week 2 (Model 2). In both models, each hypothesis was represented by a specific model parameter within a unified statistical learning framework. Hypotheses I and III correspond to fixed effect coefficients, while Hypothesis II is addressed through a variance component.

#### Model 1 - Private Information

Model 1, addressing private information acquisition, included an overall intercept, and test instance (continuous) and treatment (two levels, trained versus naive) as fixed effects. In addition, this model also included random intercepts for each individual and slopes for trained individuals. In detail this model is defined as:

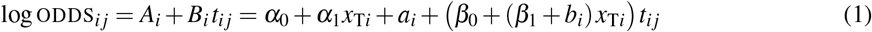

where the probability *P*_*ij*_ of success (*y*_*ij*_ = 1) is modeled via the odds ODDS_*ij*_ = *P*_*ij*_/(1 − *P*_*ij*_), allowing interpretation in terms of odds ratios. The combined, subject-specific intercept *A*_*i*_ represents the baseline odds of reaching the region of interest prior to any learning (Test Instances 1–2; Figure 3). The slope *B*_*i*_ reflects the individual learning rate, with ODDS_*i j*_ expected to increase as a function of *t*_*ij*_, the number of instances since the task was first solved (bounded by 15; Figure 3). This formulation yields a sigmoidal learning curve for *p*_*ij*_ over time (Figure 4). Training status was dummy-coded as *x*_T*i*_ = 1 for trained individuals and 0 otherwise. The learning slope *B*_*i*_ therefore consists of a baseline learning rate β_0_ for untrained individuals, a fixed-effect increase β_1_ for trained individuals, and a subject-specific random deviation *b*_*i*_. The intercept *A*_*i*_ is specified analogously. Random effects *a*_*i*_ and *b*_*i*_ are assumed to follow a bivariate normal distribution with standard deviations τ_*a*_ and τ_*b*_ and correlation ρ. The random slope *b*_*i*_ was included only for trained individuals, who were of primary interest. In this model, β_1_ > 0 corresponds to Hypothesis I (P1), indicating that individuals can learn to feed inside the cylinder, while τ_*b*_ > 0 corresponds to Hypothesis II (P2), reflecting consistent individual differences in learning rates. An alternative formulation including an indicator variable *x*_solved*ij*_, equal to 1 if the individual had solved the task prior to instance *j*, was evaluated to allow for more abrupt learning transitions but was not supported by AIC-based model selection.

**Figure 3.**
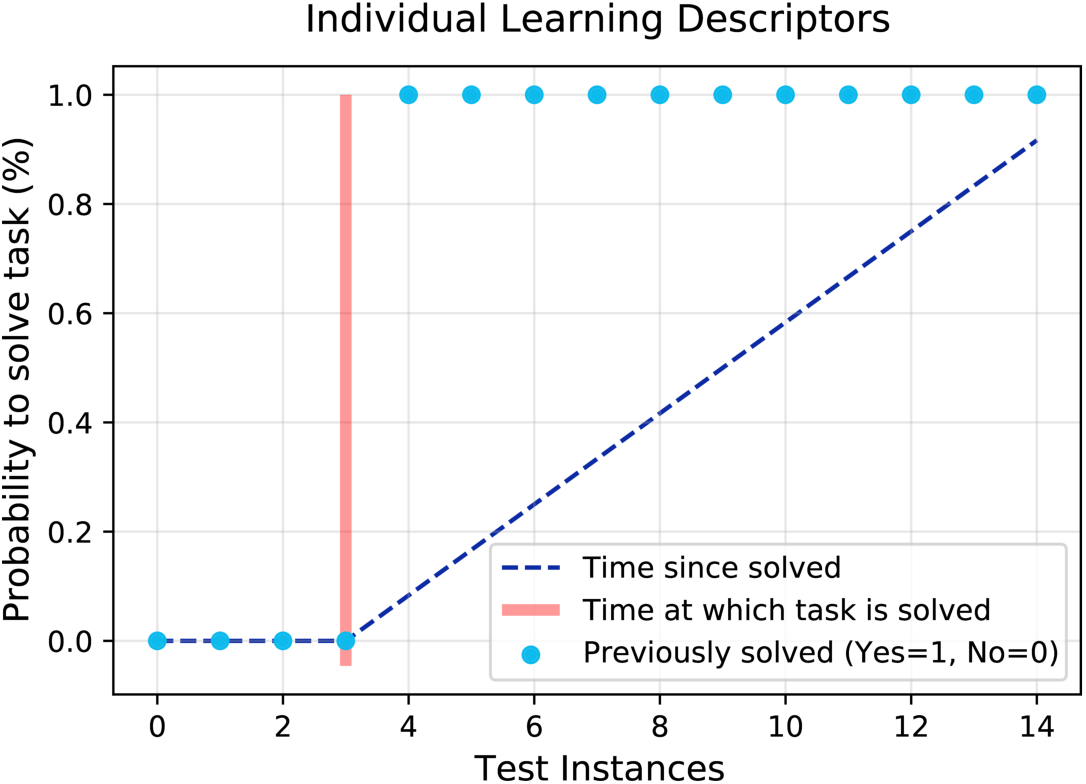
Sketch of the descriptors used to define individual learning outcome. Definition of the ‘instances since initially solved’ *t*_*ij*_ used as variable for individually describing the learning process (see Figure 4 & 5). Until the food inside of the cylinder was first found by individual *i* at test instance *J*_*i*_ = min{*j* : *y*_*ij*_ = 1}, no training effect can occur and *t*_*ij*_ = 0 for *j* < *J*_*i*_. After that, individual training commences and training time monotonically increases as *t*_*ij*_ = *j* − *J*_*i*_.

**Figure 4.**
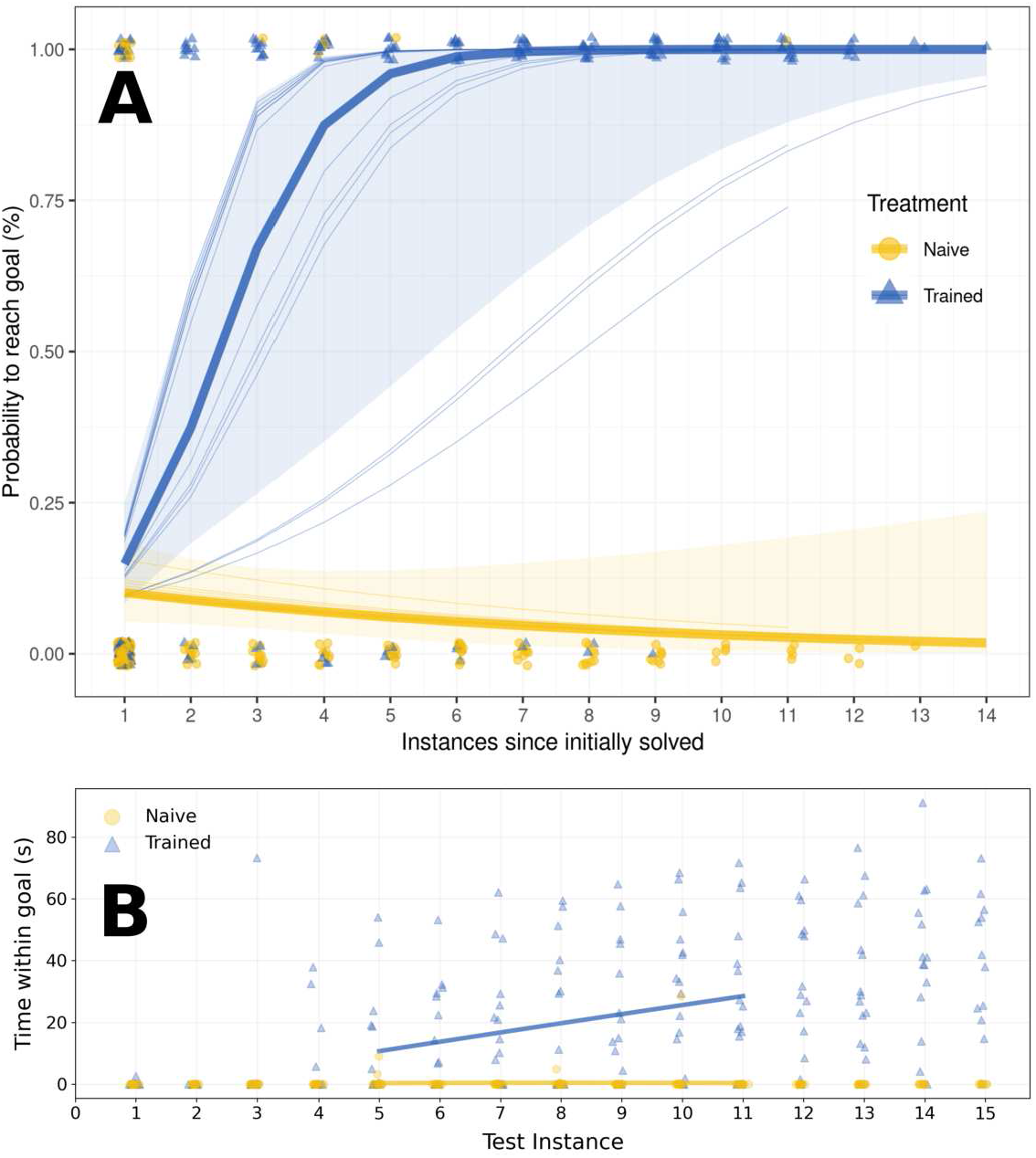
Learning outcome of the two treatment groups (trained and naive) during the first week. A: Model output shown as estimated success probability as a function of solved instances (lines, thin: individual; bold: marginal probability for trained/naive groups) and raw data (points). Instances on the x-axis are counted relative to the first time the goal was reached. Shaded areas represent 95% confidence intervals. B: Time spent within the goal area across treatment groups and over all test instances in the first week. A truncated linear fit is shown as a trend line (instances 5–11), estimated over all data points and for each treatment group. Jitter was applied along x to reduce overlap.

#### Model 2 - Social Information

Model 2, designed to compare learning dynamics among individuals with visual access to partners of different training histories, included an overall intercept, as well as test instance (continuous) and treatment (four levels: NN, NT, TT, and TN; e.g., NT denotes “naive with trained partner”) as fixed effects. In addition, this model included random intercepts for each individual and random slopes for trained individuals. In detail, the model is defined as:

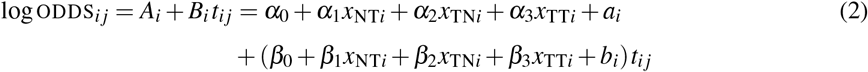

where *x*_NT*i*_ = 1 if individual *i* was naive in Week 1 but paired with a trained partner, and 0 otherwise; *x*_TN*i*_ and *x*_TT*i*_ are defined analogously. The fixed effect β_0_ represents the baseline learning rate for the reference group NN, while β_1_, β_2_, and β_3_ capture deviations associated with the remaining treatment groups. In particular, β_1_ ≠ 0 tests the first prediction of Hypothesis III (P3), indicating altered learning performance of naive individuals paired with trained partners, while the contrast between β_2_ and β_3_ tests the second prediction (P4), capturing whether experienced individuals paired with naive partners show reduced performance relative to those paired with experienced partners. The intercepts α_0_, … , α_3_ reflect the baseline success probability of (un)trained fish, potentially also allowing to capture instantaneous effects of having an (un)trained partner. Random effects were specified as in Model 1 to account for subject-specific variation.

## Results

### I. Amazon mollies are able to quickly learn the foraging task

We first examined whether clonal fish were capable of learning to feed from the provided cylinder. We addressed this question using Model 1, which captured the structure of the data well (marginal *R*^2^ = 0.83, conditional *R*^2^ = 0.839; [62]). At baseline, untrained fish exhibited odds of approximately 1:9 (corresponding to a probability of 0.10) of reaching the region of interest during a test instance. This estimate is given by the intercept *α*_0_ = −2.18 (CI = [−2.80, −1.57]) averaging over the random intercept *a*_*i*_ with estimated standard deviation τ_*a*_ = 0.19, and reflects the likelihood that a random individual enters the region of interest without prior experience (see Figure 3, Test Instances 0–2). Trained individuals, who were fed inside the region of interest, showed a slightly higher baseline probability of reaching the region of interest, with odds increased by a factor of exp(*α*_1_) = 1.55 (CI = [0.67, 3.56], *p* = 0.302). As expected from randomization, this difference was not statistically significant. For untrained individuals, we observed a weak and non-significant negative learning effect of prior cylinder entry (*β*_0_ = −0.14, CI = [−0.35, 0.065]). In contrast, trained individuals exhibited a strong and significant positive learning effect (*β*_1_ = 1.37, CI = [0.60, 2.14], *p* < 0.001). Once trained individuals had solved the task for the first time (Figure 3, Test Instances > 3), their likelihood of subsequently reaching the food source increased markedly. Specifically, for a trained individual, the odds of success more than tripled with each additional visit, yielding an odds ratio of OR = exp(*β*_0_ + *β*_1_) = 3.42 (CI = [1.60, 7.30]), under the conditional *ceteris paribus* interpretation used throughout. Figure 4 illustrates the estimated mean learning curves for trained and untrained individuals, showing the probability *P*_*ij*_ of solving the task as a function of experience (*t*_*ij*_).

### II. Clonal Amazon mollies individually differ in learning ability

We next examined whether individual variation in learning ability, specifically the probability of reaching the goal area, was detectable across individuals. Using the same model as in Section I (Model 1), we assessed individual differences via the random slope *b*_*i*_ for the learning rate of trained individuals. The estimated standard deviation of the random slopes (τ_*b*_ = 0.74) indicates substantial inter-individual variation, accounting for approximately 60% of the mean learning rate (i.e., τ_*b*_/(*β*_0_ + *β*_1_) ≈ 0.6). Testing for τ_*b*_ > 0 confirmed significant among-individual differences in learning behavior (*p* < 0.001). In addition to variation in learning rate, the standard deviation of the random intercepts (τ_*a*_ = 0.43) may reflect differences in baseline exploratory behavior, although this effect was not statistically significant (*p* = 0.386). Support for individual-level variation is further provided by model comparison: Model 1 with random effects was strongly preferred (cAIC = 259) over an identical model without random slope (cAIC = 314) or without any random effects (AIC = 342).

### III. Evident social effects of an experienced partner hinder naive learning

During the second experimental week, paired observations allowed us to assess whether learning performance differed depending on the training status of the social partner. We tested the two predictions of Hypothesis III (P3 & P4) using Model 2 (illustrated in Figure 5A): first, whether naive individuals paired with trained partners showed improved learning relative to naive-naive pairs, and second, whether experienced individuals showed reduced performance when paired with naive partners relative to trained-trained pairs. This model captured the variance in the data well (marginal *R*^2^ = 0.716, conditional *R*^2^ = 0.903). In Week 2, naive individuals paired with naive partners (reference group NN) exhibited baseline probabilities of initially entering the region of interest similar to those in Week 1 (odds exp(*α*_0_) = 0.07, CI = [0.02, 0.25]). As expected, baseline probabilities were substantially higher for experienced individuals (TN vs. NN: OR = exp(*α*_1_) = 24.74, CI = [2.82, 216.76], *p* = 0.004), consistent with the training effects observed earlier. There was no evidence that a partner’s experience increased an individual’s probability of initially entering the cylinder. Instead, we observed a tendency toward reduced baseline performance when individuals were paired with experienced partners, both for naive (NT vs. NN: OR = 0.39, CI = [0.04, 4.04], *p* = 0.432) and trained individuals (TT vs. TN: OR = 0.92, CI = [0.10, 7.79], *p* = 0.938). While these effects were not statistically significant and subject to considerable uncertainty, they suggest a decrease of approximately 61% in naive individuals and 8% in trained individuals when paired with experienced partners.

**Figure 5.**
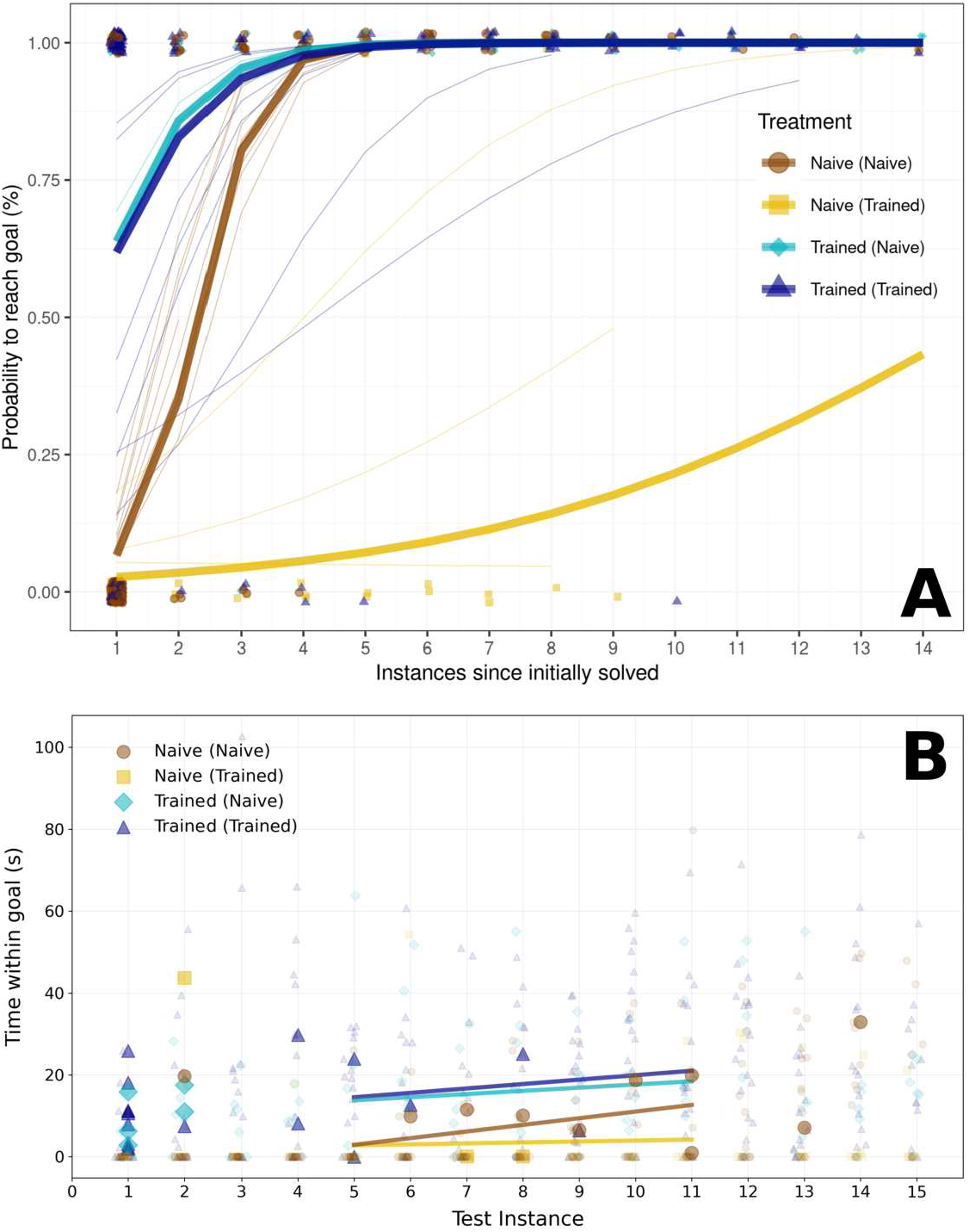
Learning outcomes of four treatment groups, depending on the focal individual and partner (in brackets): Naive (Naive): *N* = 12, Naive (Trained): *N* = 6, Trained (Naive): *N* = 6, Trained (Trained): *N* = 12. A: Model output of the second week of training in a social context, shown as estimated success probabilities (lines, thin: individual; bold: marginal probabilities averaged within each group) and raw data (points). Instances on the x-axis are counted relative to the first time the goal was reached. B: Time spent within the goal area across all treatment groups and over all test instances in the second week. First solved instances are shown with large icons and higher contrast; all remaining data are shown with reduced contrast. A truncated linear fit is shown as a trend line (instances 5–11), estimated over all data points for each treatment group. A slight jitter was applied to reduce overlap.

Crucially, a significant negative effect of partner experience emerged for the learning rate of naive individuals. Relative to the NN reference group (*β*_0_ = 2.03, CI = [1.14, 2.92]), naive individuals paired with experienced partners showed a reduced learning rate (*β*_1_ = −1.77, CI = [−2.99, −0.56], *p* = 0.004), corresponding to an odds ratio of OR = 0.17 (CI = [0.05, 0.57]) for revisiting the region of interest after the first successful entry. For experienced individuals, the negative effect of having an experienced partner was weaker and not statistically significant (TT vs. TN: OR = 0.87, CI = [0.20, 3.64], *p* = 0.847).

Overall, our results consistently indicate reduced learning performance in individuals paired with experienced partners compared to those paired with naive partners. This effect is most pronounced, and statistically significant, for the learning rate of naive individuals paired with experienced conspecifics.

## Discussion

In this study, we show that Amazon mollies can be trained in an operant conditioning task (P1), that they exhibit consistent among-individual differences in learning performance (P2), and that the presence of a task-experienced social partner can reduce learning success, particularly in task-naive individuals. Contrary to our prediction (P3) that naive fish would benefit from experienced partners, naive individuals paired with trained partners performed worse rather than better. The predicted reduction in performance of trained fish paired with naive partners (P4) was not statistically supported. We discuss potential mechanisms below.

Amazon mollies learned to associate food with a specific location within a few days and after relatively few training sessions. This finding aligns with extensive evidence from fish cognition research demonstrating that fish are capable learners with sophisticated cognitive abilities [63–66]. Similar learning capacities have been reported for both parental species of the Amazon molly, *P. latipinna* and *P. mexicana*, as well as for the closely related guppy (*P. reticulata*), all of which are capable of operant conditioning and reversal learning [30, 67]. It is therefore notable that we observed comparable cognitive abilities in the clonal Amazon molly, extending the evidence for operant learning capacity to this unstudied species.

Our results further reveal consistent individual variation in learning trajectories during the solitary phase of the experiment. While consistent individual differences in behavioral and cognitive traits are well documented [31, 32], including in clonal Amazon mollies [33, 34], learning as a varying trait among isogenic individuals has only recently been examined in detail, notably in *Drosophila melanogaster* [44]. Here, we extend this concept to a naturally occurring clonal vertebrate, demonstrating that genetically identical individuals can nonetheless differ consistently in their learning performance. Potential sources of this variability include pre-birth processes such as epigenetic differences, maternal effects [68], and developmental stochasticity [69], as well as post-birth influences including individual experience [70] and environmental conditions [33, 71]. Similar to genetic diversity within a population, such behavioral variation may reflect adaptive flexibility, allowing individuals to cope with environmental change and uncertainty [72, 73]. Although all individuals in our study were genetically identical and reared under near-identical conditions, they originated from different mothers, which may have contributed to the observed variation [34]. Identifying the relative importance of these mechanisms will require further experimental investigation.

We demonstrate that the skill level of a social partner strongly influences individual learning, but in an unexpected direction. Naive individuals paired with trained partners exhibited slower learning than naive individuals paired with other naive partners. In contrast, trained individuals did not differ significantly in learning performance when paired with naive or trained partners, although the trend suggested mildly inhibitory rather than facilitative effects of observing an experienced conspecific.

Overall, the presence of a highly skilled social partner did not enhance learning; instead, naive individuals benefited more from pairing with similarly naive partners. Building on the concept of social partners providing information of varying quality, we propose several mechanisms which may explain why naive-naive pairs outperformed naive-experienced pairs.

First, goal locations were mirrored between social partners (Figure 2), meaning a partner’s behavior did not necessarily provide transferable spatial information. Similar to Swaney et al. [13], experienced individuals in this study may have already established stereotyped task-solving routines, such as approaching the goal from a particular direction or at a particular time, that did not align with an observer’s own spatial requirements, thereby creating a mismatch between observed and required behavior. Naive individuals paired with naive partners showed rapid learning in Week 2, which is in line with the idea that two simultaneously naive individuals acquire spatially congruent information, likely producing more synchronized exploratory behavior than pairs at different skill levels. Consistent with this, Kohn [21] suggests that the variable, exploratory behavior typical of naive individuals can sustain attention and reinforce learning, whereas the repetitive, stereotyped behavior of experienced individuals may be less informative to observers.

Second, our experimental design prevented individuals from seeing others directly access or consume food. In studies reporting stimulus or local enhancement effects, observers typically witness demonstrators receiving an immediate reward [4]. The absence of visible reinforcement in our study may therefore have limited social learning, particularly from experienced partners who quickly disappeared into the cylinder while performing the task and receiving the reward. Supporting this interpretation, trained partners had little effect on naive individuals’ initial approaches to the cylinder, but strongly reduced their likelihood of revisiting the region of interest. This pattern suggests a more complex mechanism than simple spatial misguidance and argues against facilitative processes such as stimulus enhancement [74] or local enhancement [75] during learning.

While we found no evidence that trained individuals were impaired by naive partners, confidence intervals were broad and the sample size for asymmetric pairings was limited (N=6), such that weaker effects cannot be excluded. Consistent with our findings, several studies using full-contact demonstrator–observer designs in path-learning tasks have reported weak or absent positive effects of highly skilled partners. In guppies, naive individuals preferentially followed familiar but less skilled partners through unfamiliar maze configurations [13]. Similarly, in zebrafish, social information has been shown to promote food-income equality, with observers relying on visual behavioral cues from successful demonstrators to locate food [17]. In pigeons, Biederman and Vanayan [22] demonstrated that naive individuals observing demonstrators that initially performed at chance level and gradually improved subsequently outperformed those observing highly proficient demonstrators in both learning speed and overall task accuracy.

Together, these findings suggest that the value of social information is shaped not only by demonstrator proficiency but by the broader informational context in which learning occurs. Differences in information among group members are known to shape collective dynamics [76, 77], and factors such as information quality, uncertainty, and redundancy are likely to play central roles in the learning behavior of gregarious animals [1, 78, 79]. Viewed as a process of information acquisition and integration over time, learning provides a useful lens through which to investigate the temporal structure of these interactions.

## Conclusion

Although genetically identical, the individuals tested here exhibited consistent differences in learning behavior, in line with previous work showing that stable among-individual differences are common even in clonal animals [33, 34, 80]. Our study extends the operant learning framework to a naturally clonal fish species in which learning has not previously been examined, offering an interpretable approach for quantifying learning efficiency and inter-individual variation through jointly designed experiments and statistical models in which biologically meaningful parameters, such as learning rate and baseline exploration, are explicitly represented.

In summary, our findings demonstrate that the experience and prior knowledge of social partners can directly influence individual learning outcomes, and that the social context in which learning occurs shapes how effectively individuals adapt to novel situations. The counter-intuitive finding that naive individuals benefited more from similarly naive partners highlights the importance of information accessibility over demonstrator proficiency. The modeling framework introduced here, in which learning rate and baseline performance are explicitly represented as biologically meaningful parameters, provides a foundation for future work on the interplay between social structure, information flow, and learning in animal groups.

Returning to Machiavelli’s observation that people follow others’ paths even when unable to fully emulate them, we add a caveat: a model’s value depends not only on proficiency, but also on whether it remains accessible and informative to those that seek to learn from it.

## Supporting Material

All supplemental files, such as code for tracking and statistical analysis, as well as the data used in this study can be found here: https://github.com/RoboFishLab/AmazomMollyLearningMS

1. Dataset, Data_AmazonMollyLearning.csv, 2026, Author: F. Francisco
2. Statistical Code, Rstats_AmazonMollyLearning.Rmd, 2026, Authors: F. Francisco, A. Stöcker, J. Lukas
3. Video Recording Script, VideoAcquisition_AmazonMollyLearning.py, 2026, Author: F. Francisco
4. Tracking Code, TrackingCode_AmazonMollyLearning.py, 2026, Author: F. Francisco

## Author Contributions

FF, DB and PR derived the research question and experimental design. FF conducted the experiments. FF and AS conducted the statistical analysis and designed the analytical structure, while JL provided statistical insight and contributed to data curation. FF wrote the initial draft of the manuscript with input from all co-authors. All authors acknowledge no conflict of interests and have proofread the final version of the manuscript.

## Ethical Note

Animal experiments were conducted under the animal experiment number #0089/21 of the German State Office for Health and Social Affairs (LAGeSo). A total of N=36 fish were used over the course of this experiment. All animals used for this research were kept under optimal holding conditions, in order to assure healthy, natural behaviour. After the experiment, animals were released to designated holding tanks. All animal facilities and maintenance protocols were kept in accordance to the LAGeSo.

